# Influence of MUC1 on trafficking of TRPV5 and TRPV6 and *in vivo* Ca^2+^ homeostasis

**DOI:** 10.1101/2022.10.22.513333

**Authors:** Mohammad M. Al-bataineh, Carol L. Kinlough, Allison Marciszyn, Tracey Lam, Lorena Ye, Kendrah Kidd, Joseph C. Maggiore, Paul A. Poland, Anthony Bleyer, Daniel J. Bain, Thomas R. Kleyman, Rebecca P. Hughey, Evan C. Ray

## Abstract

Polymorphism of the gene encoding mucin 1 (*MUC1*) is associated with skeletal and dental phenotypes in human genomic studies. Animals lacking MUC1 exhibit mild reduction in bone density. These phenotypes could be a consequence of modulation of bodily Ca homeostasis by MUC1, as suggested by the previous observation that MUC1 enhances cell surface expression of the Ca^2+^-selective channel, TRPV5 in cultured unpolarized cells. Using biotinylation of cell-surface proteins, we asked whether MUC1 influences endocytosis of TRPV5 and another Ca^2+^-selective TRP channel, TRPV6, in cultured polarized epithelial cells. Results indicate that MUC1 reduces endocytosis of both channels, enhancing cell surface expression. Further, mice lacking MUC1 lose apical localization of TRPV5 and TRPV6 in the renal tubular and duodenal epithelium. Females, but not males, lacking MUC1 exhibit reduced blood Ca^2+^. However, mice lacking MUC1 exhibited no differences in basal urinary Ca excretion or Ca retention in response to PTH receptor signaling, suggesting compensation by transport mechanisms independent of TRPV5 and TRPV6. Finally, humans with autosomal dominant tubulointerstitial kidney disease due to frame-shift mutation of MUC1 (ADTKD-MUC1) exhibit reduced plasma Ca concentrations compared to control individuals with mutations in the gene encoding uromodulin (ADTKD-UMOD), consistent with MUC1 haploinsufficiency causing reduced bodily Ca^2+^.

## INTRODUCTION

Mucin 1 (MUC1) is a heavily glycosylated, single transmembrane protein, expressed at the apical surface of many epithelial cells. Previous studies have demonstrated that MUC1 enhances activity of the Ca^2+^-selective transporter, TRPV5, *in vitro*. In this manuscript, we ask whether MUC1 influences sub-cellular localization of TRPV5 *in vivo*, whether MUC1 influences activity of other Ca^2+^-selective transporters (TRPV6), *how* MUC1 influences cellular trafficking of TRP channels, and whether MUC1 deficiency exerts an *in vivo* influence on Ca^2+^ homeostasis.

Human observational studies suggest MUC1 could participate in Ca^2+^ homeostasis. Genetic analyses find an association between *MUC1* and numerous skeletal phenotypes including: heel bone mineral density (p = 1.4 × 10^-13^)(1), estimated bone density from heel ultrasound (p = 1 × 3.9 × 10^-11^) (2), estimated bone mineral density (2.1 × 10_-6_)(3), femoral neck bone mineral density (1.5 × 10^-4^) (4), bone mineral density in older people (4.8 × 10^-4^) (5), standing height (p = 1.5 × 10^-7^) (6), childhood height (p = 1.6 × 10^-4^) (6), and need for dentures (p = 4.2 × 10^-10^) (6). Urinary MUC1 elaboration is associated with the presence of hypercalciuria and calcium nephrolithiasis (7). Though these studies demonstrate an association between *MUC1* and traits that are influenced by Ca^2+^ homeostasis, this association does not explain how MUC1 could influence Ca^2+^ homeostasis.

One clue comes from the observation that MUC1 is expressed in several epithelia that are specialized for transcellular Ca^2+^ transport (8,9). In the kidney, MUC1 is expressed in the thick ascending limb, distal convoluted tubule (DCT), connecting tubule (CT) and collecting duct (CD) (10). The DCT and CT participate in trans-cellular Ca^2+^ absorption through the activity of transient receptor potential cation channel subfamily V members 5 and 6 (TRPV5 and TRPV6, previously referred to as ECaC1 and ECaC2, respectively). In the GI tract, TRPV6 and MUC1 are co-expressed in duodenum, ileum, and colon, where trans-cellular Ca^2+^ uptake also occurs (11,12).

MUC1 physically interacts with TRPV5, as demonstrated by the ability of these proteins to coimmunoprecipitate (7). Furthermore, MUC1 enhances cell surface expression and activity of TRPV5 in cultured fibroblasts. The ability of MUC1 to increase cell surface expression of TRPV5 is dependent upon a single, glycosylated asparagine in TRPV5 (N358Q in the human ortholog) and upon expression of galectin-3, which is suggested to physically mediate the interaction between MUC1 and TRPV5. The ability of MUC1 to enhance activity of TRPV5 depends upon both dynamin and caveolin, suggesting that MUC1 enhances cell surface expression of TRPV5 by impairing endocytosis, at least in fibroblasts.

As the Asn residue in TRPV5 that is required for MUC1-dependent up-regulation of channel activity is conserved in TRPV6, we also ask whether MUC1 influences subcellular localization of TRPV6 *in vivo*. Further, does MUC1 influence the rate of TRPV6 endocytosis in polarized epithelial cells (MDCKs)? We examine MUC1-deficient animals to determine whether they exhibit altered Ca^2+^ homeostasis. Finally, we evaluate Ca^2+^ levels in patients with *MUC1* mutation causing autosomal dominant, tubulo-interstitial kidney disease (ADTKD-MUC1) to determine whether *MUC1* mutation results in haploinsufficiency, causing reduced blood Ca^2+^ levels as compared to control patients with autosomal dominant, tubulointerstitial kidney disease due to mutation in *UMOD*, encoding uromodulin (ADTKD-UMOD).

## MATERIALS AND METHODS

### Cell culture and Endocytosis assays

Madin-Darby canine kidney (MDCK 2001) cells were obtained from Kai Simons in the European Molecular Biology Laboratory in Heidelberg, Germany. Cells were grown in DMEM-F-12 (D6421) supplemented with 5% FBS (GIBCO/Thermo Fisher Scientific, Grand Island, NY) and maintained at 37°C in 5% CO_2_. Preparation of the MDCK cell line stably transfected with human MUC1 with 22 tandem repeats (MDCK-MUC1) was previously described (13). MDCK-MUC1 cells or non-transfected MDCK cells were plated at confluency on 12-well plastic dishes and transfected with EGFP-TRPV5 or TRPV6-Flag the following day using Lipofectamine 2000 (Invitrogen Thermo Fisher). Plasmid encoding GFP-TRPV5 (rabbit) and TRPV6-Flag (human) were kindly provided by Chou-Long Huang (University of Iowa) and Ji-Bin Peng (Universtiy of Alabama at Birmingham), respectively. The TRPV6 cDNA was subsequently moved to pCDNA3 (Sigma-Aldrich). Endocytosis of proteins in MDCK cells was carried out as previously described (14). Briefly, cells were treated with membrane-impermeant sulfo-NHS-SS-biotin on ice and moved to a circulating water bath at 37°C for 0, 10 or 20 min. Cells were returned to ice and surface biotin was stripped with MESNA. Cells were scraped and extracted in detergent with protease inhibitors and phosphatase inhibitors and incubated overnight at 4°C with neutravidin-conjugated beads. Beads were washed and proteins eluted in sample buffer with Δ-mercaptoethanol at 60°C for 5 minutes before SDS-PAGE and immunoblotting for either TRPV5, TRPV6, MUC1 or endogenous podocalyxin (see Table 1). Rabbit anti-GFP antibody (Invitrogen) was used for GFP-TRPV5 (dilution 1:4000). Rabbit anti-TRPV6 antibody was from Alomone Labs (dilution 1:1000). Armenian hamster anti-MUC1 cytoplasmic tail antibody CT2 was a gift from Sandra J. Gendler at Mayo Clinic, AZ (dilution 1:1000) (15). Mouse anti-podocalyxin antibodies 3F2/D8 (cell culture supernatant) were from Developmental Studies Hybridoma Bank (dilution 1:33). Secondary HRP-tagged antibodies were from Jackson Labs (dilution 1:10,000). Blots were developed using Bio-Rad Clarity ECL reagent (3 min) and a Bio-Rad Chemidoc. Data were quantified with Quantity One 4.6.6 software.

**Table 1.** Antibodies Used.

| Experiment | Antigen | Conjugation | Animal of Origin | Source | Catalog No. | Dilution |
| --- | --- | --- | --- | --- | --- | --- |
| Immunoblotting | GFP |  | Rabbit | Invitrogen | A11122 | 1:4000 |
|  | TRPV6 |  | Rabbit | Alomone | ACC-036 | 1:1000 |
|  | MUC1 C-terminus (CT2) |  | Armenian hamster | Sandra J. Gendler (15) |  | 1:2000 |
|  | podocalyxin |  | Mouse | DSHB | 3F2/D8 | 1:33 |
| Immuno-fluorescence microscopy | TRPV5 |  | Rabbit |  |  | 1:50 |
|  | Rabbit IgG | Cy3 | Goat |  |  | 1:800 |
|  | Parvalbumin |  | Guinea Pig |  |  | 1:100 |
|  | Guinea Pig IgG | Alexa-488 | Goat |  |  | 1:400 |
|  | TRPV6 |  | Rabbit | Alomone Labs | Cat #: ACC-036 | 1:50 |

### Animal care

Mice were housed at the University of Pittsburgh Department of Laboratory Animals. Experimental procedures were approved by the University of Pittsburgh Institutional Animal Care and Use Committee. Mice were propagated in the C57Bl/6J background. All experimental mice were the product of crosses between male and female *Muc1^+/−^* mice. Genotyping was performed as previously described (16). All *Muc1^+/+^* (control) mice were littermate controls. Mice were fed Prolab Isopro RMH 3000, LabDiet chow (1.09% Ca2+, 0.24% Mg2+, 0.94% K+ and 0.23% Na+) and water purified by reverse osmosis. They were maintained on a 12hour/12hour light/dark cycle.

Urine collection was performed as follows: to ensure voiding and to prevent volume depletion, animals were first injected with 7.5% (volume / body weight) sterile normal saline solution (NSS), then and placed in a metabolic cage. Urine voided in the first 30 minutes was discarded. Urine was then collected over the next 3.5 hours. Animals were sacrificed, and bladder urine was aspirated and combined with urine collected in metabolic cages.

At time of sacrifice, mice underwent non-survival surgery under isoflurane anesthesia to collect blood, kidney, and duodenum specimens.

To examine urinary response to PTH receptor activation, mice were injected with the stable PTH analog, teriparatide. Mice were given an injection of vehicle alone (5% volume / body weight NSS, i.p.), and urine was collected in metabolic cages as six one-hour fractions. Two days later, mice were again injected with 5% body weight NSS, this time with 150 μg/kg teriparatide. Urine was again collected in metabolic cages as six one-hour fractions to assess urinary excretion of Na, phosphorus, and Ca.

### Immunofluorescence confocal microscopy

Kidney or duodenum were placed in 4% paraformaldehyde in PBS for 16 h at 4 °C. After cryoprotecting the slices by immersion in 30% sucrose in PBS-0.02% azide, 5 μm thick cryosections were prepared as described previously (17,18). Immunofluorescence labeling was subsequently performed using rabbit TRPV5 or TRPV6 antibody, followed by a secondary goat antirabbit antibody coupled to CY3 (Table 1). Immunolabeled tissues were mounted in *VECTASHIELD*^®^ mounting medium (Vector Laboratories) and imaged in a confocal laser scanning microscope (Leica TCS SP5, Model upright DM 6000S, Leica Microsystems Inc., Buffalo Grove, IL, USA) using a 63x objective with identical laser settings for all samples. Immunofluorescence images were analyzed using FIJI, by investigators blinded to genotype (19). Signal intensity was measured along the length of lines drawn from the tubular or intestinal lumen, across the cell surface, and into the cytoplasmic space (avoiding nuclei). Resulting line scans were normalized to maximum height, then averaged for comparison between genotypes using Igor Pro software (Wavemetrics, Inc.). Direct comparison of cytoplasmic ? apical staining intensity was performed in Fiji by drawing boxes of arbitrary size within the cytoplasmic space and across the region of peak staining intensity at the cell apex.

### Metabolite measurement

Whole blood electrolytes were measured using an iSTAT blood analyzer (Abbott). Plasma 1,25(OH)_2_ vitamin D was measured through the Charles and Jane Pak Center for Mineral Metabolism and Clinical Research at UT Southwestern Medical Center (Dallas, TX). PTH was measured from plasma separated from whole blood by centrifugation at 4 °C in EDTA-containing tubes, flash-frozen in liquid nitrogen, and stored at −80°C. Plasmas were thawed on ice, and PTH levels were measured in plasma diluted 1:5 using a mouse PTH ELISA (MyBioSource). Urine Na, Ca, and phosphorus were measured in samples diluted in 2% nitric acid using a Perkin Elmer NeXION 300x inductively-coupled mass spectrometer. Stool Ca^2+^ measurement was performed by ICP-MS following micro-wave assisted extraction in 6% nitric acid (Center for Applied Isotope Studies, University of GA).

### Human subjects

Human data for the current study were provided by the Wake Forest Rare Inherited Kidney Disease Registry, as approved by the Wake Forest School of Medicine Institutional Review Board, in adherence with the Declaration of Helsinki (15). *MUC1* sequencing was performed either at the Broad Institute using mass spectrometry-based probe extension (16), or at the Charles University, using Illumina and SMRT methods. (17) Plasma creatinine and total calcium levels were measured using standard clinical methods at the Wake Forest Baptist Health clinical pathology laboratory. For each individual, an average calcium and eGFR value was calculated based on the first three available measurements. Individuals were excluded if they had mean eGFR less than 60 ml/min or had nonphysiologic eGFR (greater than 200 ml/min) or Ca^2+^ (greater than 12 mg/dL or less than 8 mg/dL).

### Statistics

Statistical comparisons were performed in GraphPad Prism, version 9.4.0. Outliers were identified using the ROUT method (Q = 1%), and excluded prior to statistical analysis. Specific statistical tests used are described in figure legends.

## RESULTS

MUC1 was previously shown to enhance cell surface expression of TRPV5 in a dynamin and caveolin-1-dependent fashion in fibroblasts, consistent with an influence on channel endocytosis. We asked whether MUC1 enhances cell surface expression of channels in polarized epithelial cells, using cell surface biotinylation of proteins in Madin-Darby canine kidney (MDCK) cells with, and without, stable over-expression of MUC1. We confirmed that MUC1 enhances cell surface expression of TRPV5 in polarized epithelial cells in culture (Figure 1A). Stable expression of MUC1 in MDCK cells increased TRPV5 cell surface expression (5.29 +/− 0.71 % with MUC1 vs. 4.22 +/− 0.45 % without MUC1, N = 4 for each, p < 0.05 by Student’s t-test). The increase in cell surface expression was associated with significantly decreased rates of TRPV5 endocytosis in cells with MUC1.

**Figure 1.**
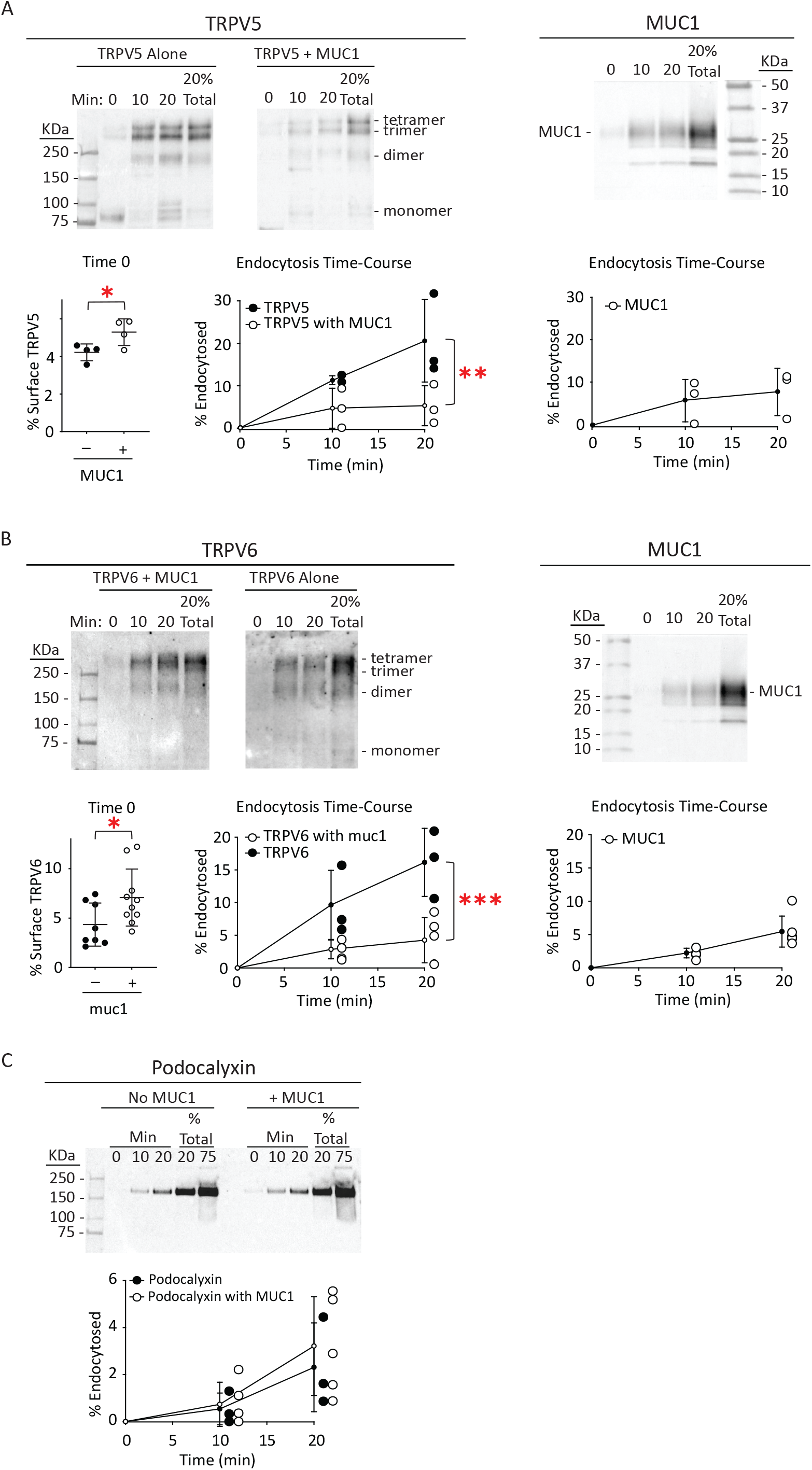
MUC1 increases cell surface expression and decreases rates of endocytosis selectively for TRPV5 and TRPV6 in polarized epithelial cells. **A.** Cell surface expression and endocytosis of TRPV5 were examined. MDCK cells with, or without, stable expression of MUC1 were transiently transfected with TRPV5-GFP. Cell surface proteins were labeled with membrane-impermeant sulfo-NHS-SS-biotin on ice and moved to a circulating water bath at 37°C for 0, 10 or 20 min. Cells were returned to ice and surface biotin was stripped with MESNA. Cell lysates were incubated overnight at 4 °C with neutravidin-conjugated beads. Beads were washed and proteins eluted in sample buffer with β-mercaptoethanol at 60°C for 5 min before SDS-PAGE and immunoblotting for either TRPV5 with anti-GFP antibodies or for MUC1 with anti-cytoplasmic tail (CT2) antibodies. Differences in oligomerization state of TRPV5-GFP were the result of the overnight incubation at 4 °C and not indicative of changes *in situ* (see Figure S-1). Therefore, all oligomerization states were included for quantification of TRPV5-GFP. Time 0 represents basal cell surface expression of TRPV5. MUC1 increased TRPV5 cell surface expression (p < 0.05 by Student’s t-test). Endocytosis time-courses (N = 3) show that MUC1 reduces TRPV5 endocytosis (p < 0.01 by two-way ANOVA). Endocytosis of TRPV5 in the presence of MUC1 did not differ significantly from that of MUC1 itself. Representative blots are shown for each analysis. **B.** Cell surface expression and endocytosis time-course of TRPV6 with, and without, co-expression with MUC1 were examined as in A. Blots were developed with rabbit anti-TRPV6 or anti-MUC1 cytoplasmic tail antibodies. Time 0 biotinylated TRPV6 demonstrated increased basal cell surface TRPV6 in the presence of MUC1 (p < 0.05 by Student’s t-test). MUC1 co-expression significantly reduced TRPV6 endocytosis (p=0.001 by two-way ANOVA). Representative blots are shown for each analysis. **C.** Endocytosis of an endogenous protein, podocalyxin, in MDCK cells is no different in the presence or absence of MUC1 expression (p = NS by two-way ANOVA). Representative blots are shown for each analysis. Error bars represent standard deviation of the mean.

The percentage of TRPV6 at the cell surface in the presence of MUC1 (7.08 +/− 2.89 %, N = 10) was also significantly different than in the absence of MUC1 (4.34 +/− 2.18 %, N = 8, p < 0.05 by Student’s t-test) (Figure 1B). As with TRPV5, MUC1 significantly retarded endocytosis of TRPV6. In the presence of MUC1, endocytosis of these channels resembles that of MUC1 itself. In contrast, MUC1 co-expression does not alter endocytosis rate of the endogenous cell surface protein podocalyxin (Figure 1C).

Having confirmed that MUC1 slows endocytosis and enhances cell surface expression of both TRPV5 and TRPV6 in polarized cells in culture, we then asked whether MUC1 influences sub-cellular localization of these channels *in vivo*. Kidney and duodenum sections from *Muc1^-/-^* or from *Muc1^+/+^* littermate control mice were immunostained for TRPV5 and TRPV6 (Figure 2). In kidney, both TRPV5 and TRPV6 are redistributed from the cell apex toward the cytoplasm in *Muc1^-/-^* mice (p < 0.0001 by Student’s T-test for cytoplasm/apical staining for both TRPV5 and TRPV6 in kidney). Similarly, in duodenum, apical localization of TRPV6 is lost in *Muc1^-/-^* animals (p < 0.0001 by Student’s T-test for cytoplasm/apical staining).

**Figure 2.**
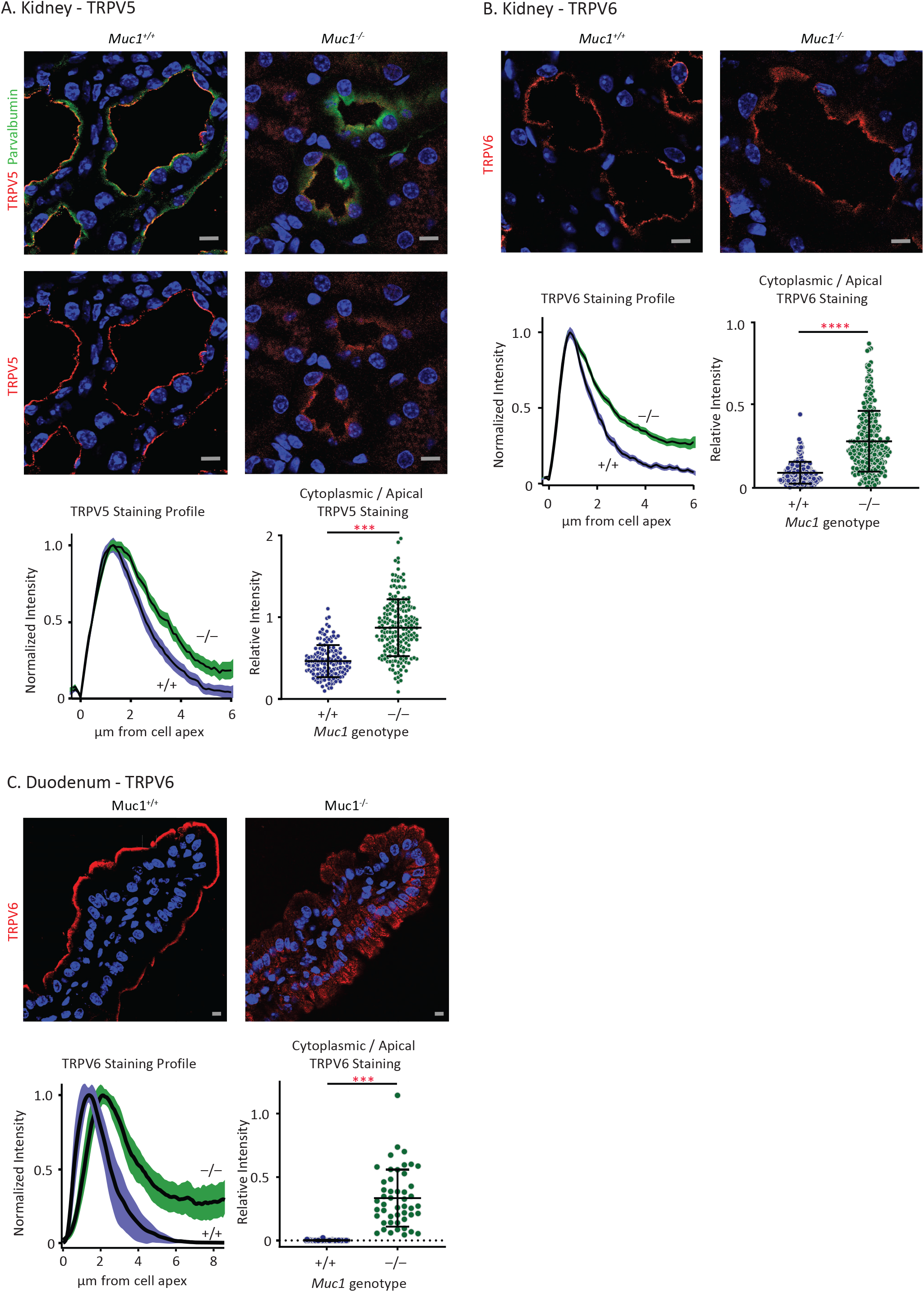
Absence of MUC1 *in vivo* shifts localization of TRPV5 and TRPV6 away from the cell apex. **A.** Representative immunofluorescence images show redistribution of TRPV5 from the renal tubule cell apex toward cytoplasm in kidney from *Muc1^-/-^* mice. Red represents TRPV5 staining. Green represents parvalbumin staining, indicating early distal convoluted tubule. Blue represents TO-PRO-3 staining. Panel below shows the mean of line scans of TRPV5 staining intensity from the cell surface, extending into the cytosol from *Muc1^-/-^* mice and from *Muc1^+/+^* controls (one linescan per TRPV5 expressing cell). Y-axis represents mean normalized intensity in early DCT epithelial cells. X-axis indicates distance from apparent cell surface. Shading behind black line represents 95% confidence interval of line scans (N = 3 animals per genotype; n = 251 cells for *Muc1^+/+^* mice; n = 332 cells for *Muc1^-/-^* mice). Note divergence of linescans within the cytoplasmic region, indicating more TRPV5 staining in the cytosol in *Muc1^-/-^* mice. Second panel represents mean cytoplasm/apical staining intensity, as calculated from arbitrarily chosen cytosolic and apical cell regions. Cytoplasm/cell apex staining of TRPV5 is greater in *Muc1^-/-^* mice (N = 3 animals per genotype, n = 147 cells for *Muc1^+/+^* mice; n = 192 cells for *Muc1^-/-^* mice; p < 0.0001 by Student’s T-test). **B.** TRPV6 subcellular localization in mouse kidney. Red represents TRPV6 staining. Panel below shows the mean of line scans of TRPV6 staining intensity. TRPV6 staining intensity is shifted away from the cell apex, toward cytoplasm (N = 3 animals per genotype, n = 180 cells for *Muc1^+/+^* mice; n = 380 cells for *Muc1^-/-^* mice). Adjacent panel shows cytoplasm / apical TRPV6 staining in tubular epithelial cells. Cytoplasm / apical TRPV6 staining is greater in *Muc1^-/-^* mice (N = 3 animals per genotype, n = 184 cells for *Muc1^+/+^* mice; n = 270 cells for *Muc1^-/-^* mice; p < 0.0001 by Student’s T-test). **C.** Duodenal epithelium TRPV6 staining is also shifted toward cytoplasm in *Muc1^-/-^* mice. Linescans and cytoplasm / apical staining were examined from 5 to 10 cells per villus in 9 to 10 individual villi per duodenum from a total of 3 mice from each genotype (n = 45 cells for *Muc1^+/+^* mice; n = 48 for *Muc1^-/-^* mice). Linescans and cytoplasm / apical staining both show a shift of TRPV6 staining away from cell apex, toward the cytoplasm in *Muc1^-/-^* mice (p < 0.0001 by Student’s T-test). In all images, gray scale bars represent 10 μm. Error bars represent standard deviation of the mean.

TRPV5 and TRPV6 knock-out animals exhibit systemic Ca^2+^-depletion as a consequence of impaired transcellular Ca^2+^ transport in the intestine and kidney tubule (20,21). We asked whether *Muc1^-/-^* mice exhibit reduced whole blood concentrations of Ca^2+^ and other electrolytes (Figure 3, Table 2). Blood Ca^2+^ levels were significantly lower in female *Muc1^-/-^* mice (1.17 ± 0.06, N = 6) compared to *Muc1^+/+^* females (1.23 ± 0.03, N = 10; p < 0.05). Male mice had similar blood Ca^2+^ in *Muc1^+7+^* (1.20 ± 0.03, N = 13) versus Muc1^-/-^ animals (1.20 ± 0.06, N = 12; p = NS).

**Figure 3.**
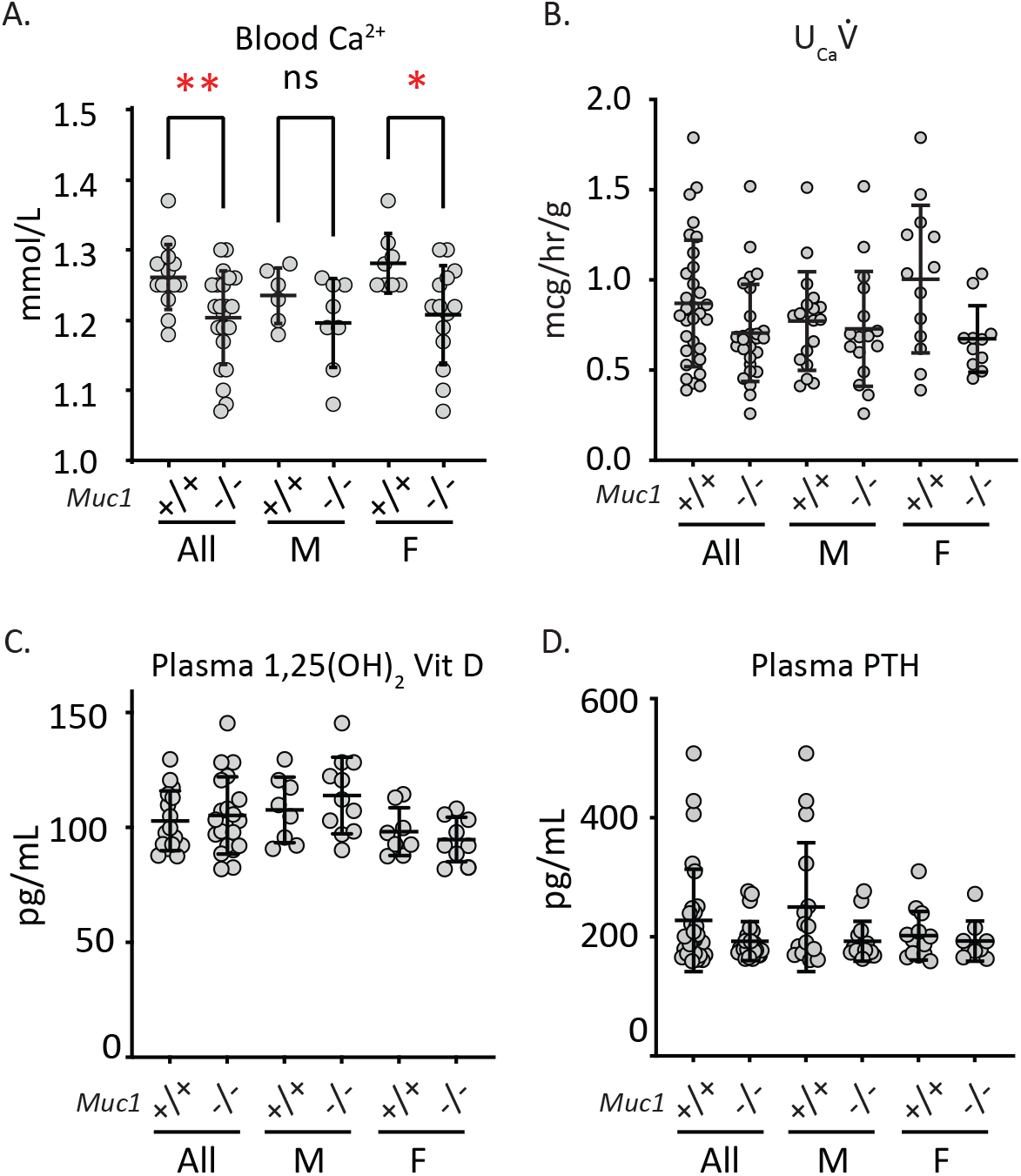
*Muc1^-/-^* mice exhibit lower blood Ca^2+^, but no differences in Ca^2+^ excretion. **A.** Blood Ca^2+^ levels are lower in female but not male *Muc1^-/-^* mice. Female *Muc1^-/-^* mice exhibited lower blood Ca^2+^ than MUC1^+/+^ controls (p < 0.05 by Student’s T-test). Male mice had similar blood Ca^2+^ in MUC1^+/+^ controls compared to Muc1^/_^ animals (p = NS). There was no genotypespecific difference when male and female mice were pooled (p = NS). **B.** Urine Ca excretion rate 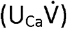 was not different in *Muc1^-/-^* mice compared to littermate controls. Urine was collected over 3.5 hours in metabolic cages, and urinary Ca excretion 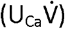 was measured. No difference in 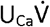 was observed overall (p = NS by Student’s T-test), or when mice were stratified on the basis of sex (p = NS in pairwise comparison by Student’s T-test). **C.** Plasma levels of 1,25(OH)_2_ Vitamin D did not differ in *Muc1^+/+^* vs. *Muc1^-/-^* mice, either overall or after stratification by sex (p = NS in pairwise comparison by Student’s T-test). **D.** PTH levels were not different in *Muc1^+/+^* vs. *Muc1^-/-^* mice either overall or after stratification by sex (p = NS in pairwise comparison by Student’s T-test). Error bars represent standard deviation of the mean.

**Table 2.** Blood Electrolytes.

| Sex<br>Genotype | All |  | Male |  | Female |  |
| --- | --- | --- | --- | --- | --- | --- |
|  | +/+ | -/- | +/+ | -/- | +/+ | -/- |
| Na<sup>+</sup> | 145 ± 1 (23) | 145 ± 2 (17) | 146 ± 1 (14) | 145 ± 2 (11) | 145 ± 2 (9) | 145 ± 2 (6) |
| K<sup>+</sup> | 4.6 ± 0.6 (23) | 4.6 ± 0.2 (17) | 4.7 ± 0.5 (14) | 4.7 ± 0.2 (11) | 4.5 ± 0.6 (9) | 4.6 ± 0.3 (6) |
| Cl<sup>-</sup> | 112 ± 2 (17) | 112 ± 2 (17) | 112 ± 3 (8) | 111 ± 2 (11) | 113 ± 2 (9) | 113 ± 2 (6) |
| BUN | 28 ± 4 (17) | 27 ± 5 (17) | 28 ± 3 (8) | 27 ± 6 (11) | 29 ± 4 (9) | 26 ± 5 (6) |

Cells show mean values (N). Na, K, and Cl units are in mmol/L. BUN: blood urea nitrogen, in mg/dL. No pairwise *Muc1*<sup>+/+</sup> vs. *Muc1*<sup>-/-</sup> comparisons were statistically significant by Student's T-test.

To examine whether MUC1-deficiency causes urinary wasting of Ca, we measured urinary Ca excretion 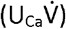 of mice in metabolic cages (Figure 3). Over 3.5 hours, MUC1^+/+^ mice excreted 0.87 ± 0.35 μg/hr/g body weight (N = 31) compared to 0.71 ± 0.27 (N = 27) in *Muc1^-/-^* mice (p = NS). No differences were observed when mice were stratified on the basis of sex: male wild-type littermates excreted 0.77 ± 0.27 (N = 18) and *Muc1^-/-^* males excreted 0.73 ± 0.32 (N = 16, p = NS); female wild-type littermates excreted 1.0 ± 0.41 (N = 13) and female *Muc1^-/-^* mice excreted 0.67 ± 0.18 (N = 11, p = NS) μg/hr/g body weight. There were also no differences in urinary Na or K excretion (not shown).

To examine whether *Muc1^-/-^* mice compensated for reduced transcellular transport by up-regulating active vitamin D or PTH production, plasma 1,25(OH)_2_ vitamin D and PTH were measured in *Muc1^-/-^* mice and in *Muc1^+/+^* littermates. Plasma levels of 1,25(OH)_2_ Vitamin D did not differ in *Muc1^+/+^* (103 ± 13 pg/mL, N = 16) vs. *Muc1^-/-^* mice (105 ± 17, N = 20; p = NS). No differences were observed after stratification of mice by sex. In males, wild-type littermates exhibited plasma 1,25(OH)_2_ Vitamin D of 108 ± 14 pg/mL (N = 8) compared to 114 ± 17 (N = 11; p = NS). In females, levels in wild-type littermates were 98 ± 11 (N = 8) compared to 95 ± 10 pg/mL (N = 9; p = NS). Nor were PTH levels different in *Muc1^-/-^* versus littermate control mice. Overall, PTH levels were 228 ± 86 pg/mL in littermate control animals (N = 30) and 193 ± 33 in Muc1^-/-^ animals (N = 24, p = NS). In males, these were 250 ± 108 (N = 16) in littermates and 193 ± 33 (N = 15, p = NS) in Muc1^-/-^ mice. In females, these were 202 ± 41 (N = 14) and 193 ± 34 (N = 9, p = NS) in littermates and Muc1^-/-^ mice, respectively.

PTH up-regulates TRPV5 cell surface expression by reducing caveolin-1-mediated endocytosis of the channel (22). Because MUC1 modulates endocytosis of TRPV5 and TRPV6 in cultured cells, responsiveness of *Muc1^-/-^* mice to a stable PTH analog (teriparatide, TPT) as compared to vehicle (normal saline, NSS) was examined (Figure 4.) In both genotypes, TPT accelerated urinary phosphorus excretion 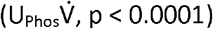 while delaying urinary Ca excretion 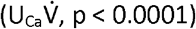 compared to excretion rates following NSS alone. The genotype-treatment interaction term was not significant by mixed effects modeling (p = NS). TPT acutely increases blood Ca^2+^, predominantly by increasing bone turn-over (23). Differences in blood Ca^2+^ could influence Ca^2+^ filtration and 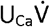. Additional mice were treated with TPT and sacrificed to confirm that TPT had similar effects on blood Ca^2+^ in *Muc1^+/+^* and *Muc1^-/-^* mice. Blood Ca^2+^ was similar in TPT-treated *Muc1^+/+^* (1.35 ± 0.05 mmol/L; N = 19) and *Muc1^-/-^* (1.38 ± 0.06 mmol/L; N = 12; p = NS) mice.

**Figure 4.**
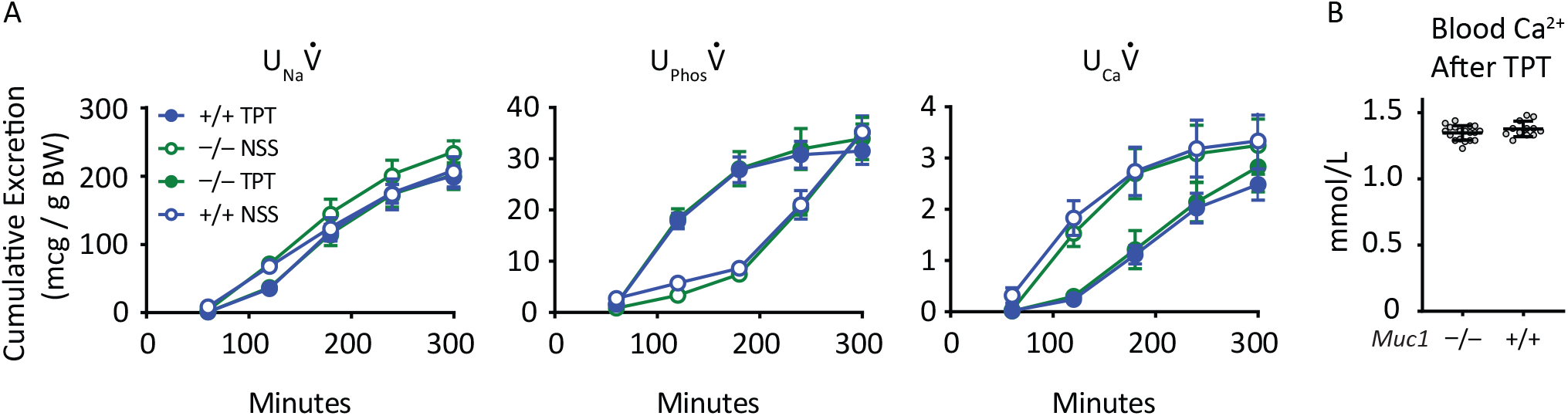
Urinary electrolyte excretion in response to the stable PTH analog, teriparatide (TPT) did not differ between *Muc1^+/+^* and *Muc1^-/-^* mice. A. Urine Na excretion 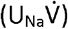, phosphorus excretion 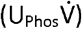, and Ca excretion 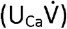 were compared following injection of 5% volume / body weight of NSS. Two days later, the same animals received 5% volume / body weight of NSS with 150 μg/kg TPT. N = 8 *Muc1^-/-^* mice and 10 MUC1^+/+^ mice. Open or closed symbols represent excretion following injection with NSS or TPT, respectively. Blue or green lines and symbols represent excretion from *Muc1^+/+^* or *Muc1^-/-^* mice, respectively. TPT influenced 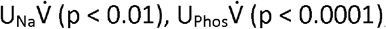, and 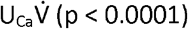, however the genotype-treatment interaction term was not significant by mixed effects modeling (p = NS). Error bars represent standard error. B. TPT induced a similar degree of hypercalcemia in Muc1^/_^ and MUC1^+/+^ animals. Blood Ca^2+^ was similar in TPT-treated *Muc1^+/+^* and *Muc1^-/-^* mice (p = NS). Error bars represent standard deviation of the mean.

To examine whether MUC1 influences Ca homeostasis in humans, plasma Ca was assessed in individuals with one working copy of *Muc1*. Autosomal dominant tubulointerstitial kidney disease can occur as a consequence of a frame-shift mutation in *Muc1* (ADTKD-MUC1) or mutation in the gene encoding uromodulin (*UMOD*, ADTKD-UMOD) (24). Both result in slow progression of interstitial fibrosis, with indolent loss of glomerular filtration with mean age of end stage kidney disease occurring in the 5^th^ decade of life. Plasma total Ca levels (ionized plus non-ionized, as is commonly measured in clinical laboratories) were compared from individuals genetically determined to have either ADTKD-MUC1 or ADTKD-UMOD. Only individuals with a glomerular filtration rate of at least 60 ml/min were examined. Overall, plasma Ca was lower in individuals with ADTKD-MUC1 than with ADTKD-UMOD (Figure 5). ADTKD-UMOD patients exhibited Ca of 9.58 ± 0.38 mg/dL (N = 72), compared to 9.41 ± 0.40 in ADTKD-MUC1 patients (N = 73; p < 0.01 by Student’s T-test). After stratification on the basis of sex, lower serum Ca^2+^ concentrations remained significant in women but not men. Men with ADTKD-UMOD exhibited Ca of 9.70 ± 0.43 (N = 27), compared to men with ADTKD-MUC1, who had Ca of 9.50 ± 0.40 (N = 33; p = NS). In women, those with ADTKD-UMOD had Ca of 9.51 ± 0.32 (N = 45) as compared to 9.34 ± 0.40 in those with ADTKD-MUC1 (N = 40; p < 0.05).

**Figure 5.**
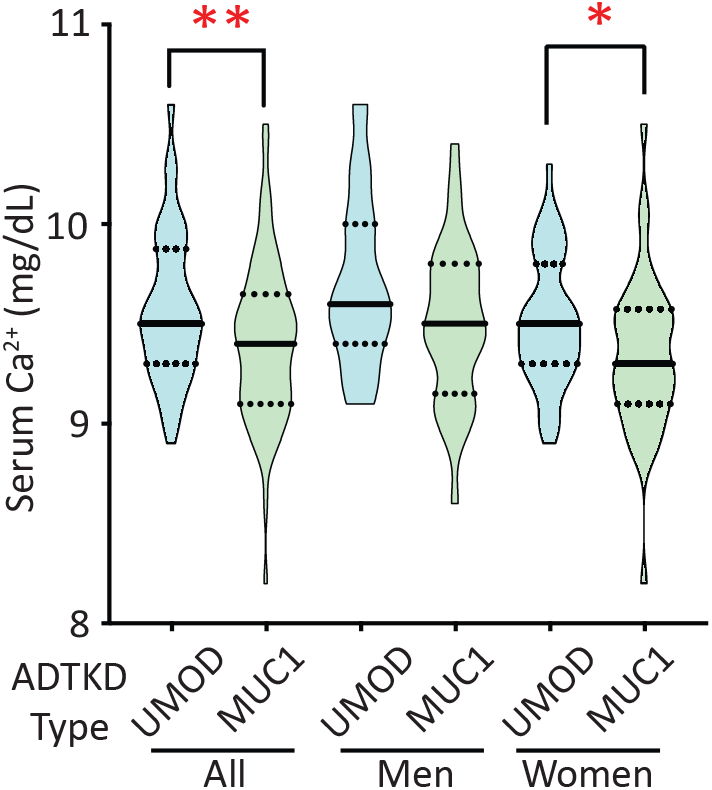
Plasma Ca is lower in individuals with autosomal dominant tubulointerstitial kidney disease due to *Muc1* mutation (ADTKD-MUC1) than in control individuals with autosomal dominant tubulointerstitial kidney disease due to *UMOD* mutation (ADTKD-UMOD). Violin plots representing plasma Ca extend from minimum to maximum values. Solid lines represent the median; dashed lines represent quartiles. Plasma Ca levels were lower in ADTKD-MUC1 patients than in controls (p < 0.01 by Student’s T-test). After stratification by sex, this difference persisted in women (p < 0.05 by Student’s T-test) but not in men (p = NS).

All data are included in the manuscript.

## DISCUSSION

Findings presented above add to evidence demonstrating a role for MUC1 in Ca ^2+^ homeostasis. 1) *Muc1^-/-^* mice exhibit reduced apical localization of both TRPV5 and TRPV6, which are critical for transcellular Ca^2+^ absorption in renal tubules and intestines. 2) *Muc1^-/-^* mice exhibit lower blood Ca^2+^. Interestingly, *Muc1^-/-^* mice exhibited no difference in urinary Ca^2+^ excretion. This could reflect a state of Ca homeostasis, albeit with lower systemic Ca stores. 3) A previous study found mild reduction in bone density in *Muc1^-/-^* mice (25). Osteoblast and osteoclast function appeared normal in *Muc1^-/-^* mice, and no compelling mechanism for the bone phenotype was described. Altered Ca^2+^ handling was not examined, but provides a good explanation for the observed bone phenotype. 4) A common *MUC1* polymorphism (rs4072037) is associated with altered bone density in humans (26,27). 5) Patients with ADTKD-MUC1, who have a frame-shift-inducing mutation in *MUC1*, exhibit lower levels of circulating Ca^2+^than individuals with ADTKD-UMOD. This is somewhat surprising, given that UMOD contributes to mineral transport in the thick ascending limb, including through support of NKCC2 activity (28). Because pharmacologic impairment of NKCC2 activity induces urinary Ca^2+^ wasting (29,30), reduced NKCC2 activity in ADTKD-UMOD might be expected to reduce blood Ca^2+^. The observation that blood Ca^2+^ levels in ADTKD-MUC1 are lower is consistent with Ca^2+^ depletion through reduced cell surface localization of Ca^2+^-selective channels, though other mechanisms could contribute. The difference in blood Ca^2+^ observed in these patients is mild, and hypocalcemia is not a recognized feature of ADTKD-MUC1. No published data address whether patients with ADTKD-MUC1 are at increased risk of bone disease associated with their chronic kidney disease.

Given the robust difference in TRP channel localization observed *in vivo* in kidney and duodenum, it is surprising that the measured parameters of Ca homeostasis – blood Ca^2+^, urine Ca excretion, PTH, and 1,25 (OH)2 vitamin D, are not more dramatically altered. Even the decrement in bone mineralization previously observed in *Muc1^-/-^* mice is relatively minor compared to the decrease in bone density observed in TRPV5^-/-^ or TRPV6^-/-^mice (20,21,25). All this suggests compensatory measures independent of these Ca^2+^-selective channels.

These data support a model in which transcellular Ca^2+^ transport is modulated by an extracellular protein lattice including MUC1, galectins, and ion channels that promotes cell surface expression of ion channels by opposing endocytosis. MUC1 enhances cell surface expression of TRPV5 in polarized epithelial cells in culture and increases apical expression of TRPV5 *in vivo*, consistent with previous findings that MUC1 promotes cell surface expression of the channel in fibroblasts (7). TRPV5 and its more broadly expressed homolog, TRPV6, share a conserved extracellular Asn residue (N438, in human TRPV5) that is N-glycosylated and is necessary for binding to galectin-3. TRPV5 exhibits galectin-binding selectivity, interacting with galectin-3, but not galectin-1. TRPV6 also depends upon MUC1 *in vivo* for apical expression. Galectin-3, through its pentameric C-terminal carbohydrate recognition domains (CRD), binds to extracellular glycans on numerous extracellular proteins (31). Among these are MUC1 and MUC2 (32). For its part, MUC1 function is also selective. It binds preferentially with galectin-3 and galectin-9 but not galectin-1, −4, −7, or −8. (33). This specificity is underscored by the finding that MUC1 selectively reduces endocytosis of TRPV5 and TRPV6, while exerting no influence on podocalyxin. Podocalyxin (gp135) is a cell-surface protein that interacts with galectin 8, but not galectin-1, −3, or −9 (34). Like MUC1, uromodulin promotes cell surface expression of TRPV5 by opposing channel endocytosis, through mechanisms that may resemble those seen for MUC1 and TRPV5 (35). All of these observations suggest that an extracellular lattice including MUC1, galectin-3, and TRPV5 and TRPV6 contributes to transcellular calcium transport.

The importance of this network in mammalian physiology is suggested by the observation that multiple genetic components originated in concert as mammals diverged from their ancestors. Mucins originated in early metazoans (36), but MUC1 arose in the earliest mammals from a precursor similar to MUC5 (37). TRPV5 and V6 arose when a common ancestral gene underwent duplication, also with the earliest mammals (38). Deletion of either *Trpv5* or *Trpv6* in mice leads to impaired Ca^2+^ handling, rarefaction of bone, and compensatory increases in 1,25(OH)_2_ vitamin D production (20,21). Thus, it appears that these proteins play an important role in mammalian Ca physiology.

In what ways this extracellular glycoprotein network may be regulated to maintain Ca^2+^ balance remains an interesting question. PTH enhances cell surface expression of TRPV5 by inhibiting its caveolin-1-mediated endocytosis (22). Because MUC1 also influences endocytosis of the channel, we predicted that *Mud1^-/-^* mice would exhibit diminished Ca^2+^ retention in response to PTH receptor activation. However, this was not observed, perhaps because other Ca^2+^ handling effects, such as paracellular Ca^2+^ reabsorption, predominated over the short course of this experiment.

In summary, the mucin MUC1 reduces endocytosis of the Ca-selective TRP channels TRPV5 and TRPV6 and promotes cell-surface localization of these channels *in vivo*, contributing positively to Ca^2+^ homeostasis.

---

**Figure S-1.**
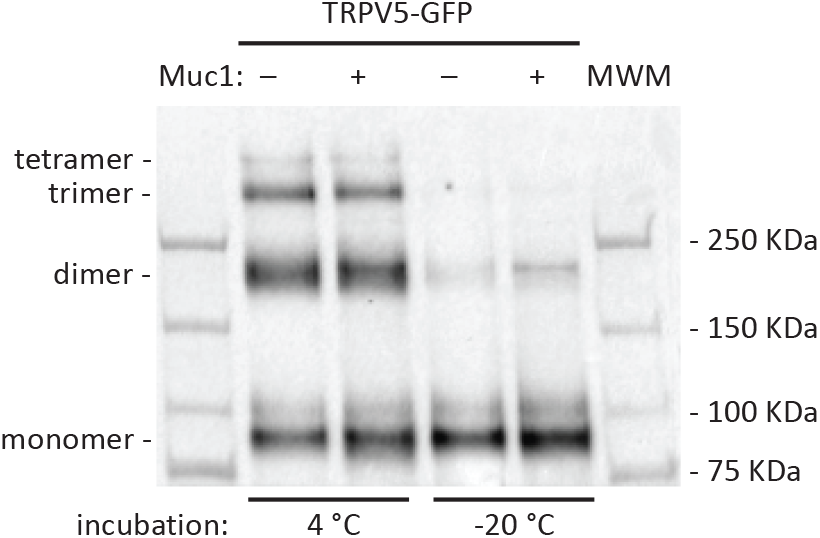
Incubation of cell lysates overnight with neutravidin beads results in oligomerization of TRPV5-GFP. A control experiment is shown comparing immunoblot of cell extract with anti-GFP antibody after incubation overnight at 4°C vs. −20°C, demonstrating that differences in stoichiometry were the result of the overnight incubation and not indicative of differences in stoichiometry in cells. Therefore, all stoichiometric bands were included for quantification of TRPV5-GFP in Figure 1.

## DECLARATION OF FUNDING

This work was supported by National Institutes of Health [K08DK110332 and T32DK061296 to E.C.R., K01DK109038 and R03DK131093 to M.M.A., R01DK038470 to R.P.H. and T.R.K] and American Society of Nephrology Carl W. Gottschalk Research Scholar Grant to E.C.R.

## CONFLICT OF INTEREST

The authors declare that they have no conflicts of interest with the contents of this article.

## Notes

### Competing Interest Statement

The authors have declared no competing interest.

